# Salvaging complete and high-quality genomes of novel microbial species from a meromictic lake using a workflow combining long- and short-read sequencing platforms

**DOI:** 10.1101/2021.05.07.443067

**Authors:** Yu-Hsiang Chen, Pei-Wen Chiang, Denis Yu Rogozin, Andrey G. Degermendzhy, Hsiu-Hui Chiu, Sen-Lin Tang

## Abstract

**Background:** Most of Earth’s bacteria have yet to be cultivated. The metabolic and functional potentials of these uncultivated microorganisms thus remain mysterious, and the metagenome-assembled genome (MAG) approach is the most robust method for uncovering these potentials. However, MAGs discovered by conventional metagenomic assembly and binning methods are usually highly fragmented genomes with heterogeneous sequence contamination, and this affects the accuracy and sensitivity of genomic analyses. Though the maturation of long-read sequencing technologies provides a good opportunity to fix the problem of highly fragmented MAGs as mentioned above, the method’s error-prone nature causes severe problems of long-read-alone metagenomics. Hence, methods are urgently needed to retrieve MAGs by a combination of both long- and short-read technologies to advance genome-centric metagenomics.

**Results:** In this study, we combined Illumina and Nanopore data to develop a new workflow to reconstruct 233 MAGs—six novel bacterial orders, 20 families, 66 genera, and 154 species—from Lake Shunet, a secluded meromictic lake in Siberia. Those new MAGs were underrepresented or undetectable in other MAGs studies using metagenomes from human or other common organisms or habitats. Using this newly developed workflow and strategy, the average N50 of reconstructed MAGs greatly increased 10–40-fold compared to when the conventional Illumina assembly and binning method were used. More importantly, six complete MAGs were recovered from our datasets, five of which belong to novel species. We used these as examples to demonstrate many novel and intriguing genomic characteristics discovered in these newly complete genomes and proved the importance of high-quality complete MAGs in microbial genomics and metagenomics studies.

**Conclusions:** The results show that it is feasible to apply our workflow with a few additional long reads to recover numerous complete and high-quality MAGs from short-read metagenomes of high microbial diversity environment samples. The unique features we identified from five complete genomes highlight the robustness of this method in genome-centric metagenomic research. The recovery of 154 novel species MAGs from a rarely explored lake greatly expands the current bacterial genome encyclopedia and broadens our knowledge by adding new genomic characteristics of bacteria. It demonstrates a strong need to recover MAGs from diverse unexplored habitats in the search for microbial dark matter.

## Background

Rapid developments in bioinformatics and sequencing methods enable us to reconstruct genomes directly from environmental samples using a culture-independent whole-genome-shotgun metagenomic approach. These genomes, also called metagenome-assembled genomes (MAGs), have become a crucial information source to explore metabolic and functional potentials of uncultivated microorganisms [1-4]. Mining MAGs quickly expands our knowledge of microbial genome, diversity, phylogeny, evolution, and taxonomy [1-4]. For example, 18,365 MAGs were identified out of a total of 410,784 microorganisms in the Genomes OnLine Database (GOLD) [5]. A total of 52,515 MAGs were assembled from diverse habitats, and the MAG collection contains 12,556 potentially novel species and expands the known phylogenetic diversity in bacterial and archaeal domains by 44% [3].

Although genome-resolved metagenomics has revolutionized research in microbiology, significant challenges need to be overcome to make MAGs more accurate, reliable, and informative [1]. First, most MAGs are derived from the metagenomic assembly of short reads [1, 6], and these short-read-derived MAGs usually comprise numerous short contigs rather than complete or nearly-complete genomic sequences, and thus important information on genomic characters is missed, such as operons, gene order, gene synteny, and promoter/regulatory regions. As of March 2021, only 177 out of 84,768 MAGs released in NCBI were complete. Second, fragmented MAGs usually miss some gene sequences and comprise unknown contaminant sequences, mistakenly assembled into the contigs [1]. Hence, low contiguity, high fragmentation, and unwanted contamination in short-read MAGs greatly affect further analyses in a variety of microbial genome-related studies.

The emergence of long-read sequencing platforms (also called third-generation sequencing platforms) such as Nanopore and PacBio provides an opportunity to improve the contiguity of MAGs and even reconstruct complete MAGs from extremely complex microbial communities [7, 8]. Recently, researchers started to develop new assemblers to reconstruct microbial genomes with high accuracy and long contiguous fragments from long-read metagenomic datasets. In 2019, Nagarajan *et al*. developed a hybrid assembler called OPERA-MS [9]. The assembler yielded MAGs with 200 times higher contiguity than short-read assemblers used on human gut microbiomes. In October 2020, Pevzner *et al*. developed metaFlye, a long-read metagenome assembler that can produce highly accurate assemblies (approximately 99% assembly accuracy) [10, 11]. The success of these newly developed assemblers becomes an important stepping-stone for reconstruction of complete MAGs with high accuracy. However, there is still much room to improve the procedures around data processing and assembling MAGs with long reads. The current study presents a new workflow for this purpose.

Our workflow combines Illumina sequencing reads and Nanopore long sequencing reads to recover many novel high-quality and high-contiguity prokaryotic MAGs from Lake Shunet, southern Siberia, one of only four meromictic lakes in all of Siberia. The lake contains stratified water layers, including a mixolimnion layer at 3.0 m, chemocline at 5.0 m, and monimolimnion at 5.5 m. From our previous 16 rRNA amplicon survey, we know that the lake comprises at least hundreds of unknown bacteria and archaea [12], highlighting the importance of mining microbial MAGs from this rarely explored lake. However, though we attempted to recover MAGs from these layers using deep Illumina sequencing with approximately 150 Gb, only one high-quality but still fragmented MAG was obtained [12]. Hence, in this study, we developed a new workflow combining Illumina and Nanopore sequencing reads by integrating several cutting-edge bioinformatics tools to recover and reconstruct MAGs with high contiguity and accuracy. We demonstrate that our newly built workflow can be used to reconstruct hundreds of complete high-quality MAGs from environmental samples in a high-complexity microbial community.

## Results and Discussions

### Reconstruction of metagenome-assembled genomes with high contiguity from Lake Shunet by combining Nanopore and Illumina sequences

To recover novel metagenome-assembled genomes (MAGs) with high contiguity without compromising accuracy, 3.0-, 5.0-, and 5.5-m deep Lake Shunet samples were sequenced by Nanopore machines individually, and the resulting long reads (LRs) were analyzed together with short reads (SRs) using a workflow we developed for this study (Fig. 1a). Originally, we only used metaFlye, a state-of-art long-read metagenome assembler that can provide 99% accuracy [10, 11], to assemble the LRs. However, recent studies found that assemblies from long reads contain numerous in-del errors, leading to erroneous predictions of open reading frames and biosynthetic gene clusters [1, 10]. Incorrectly predicting open reading frames also affects the estimation of genome completeness by single-copy marker gene method, such as checkM [13]. Hence, we used SRs from Illumina sequencing to correct the contigs generated by LRs.

**Figure 1.**
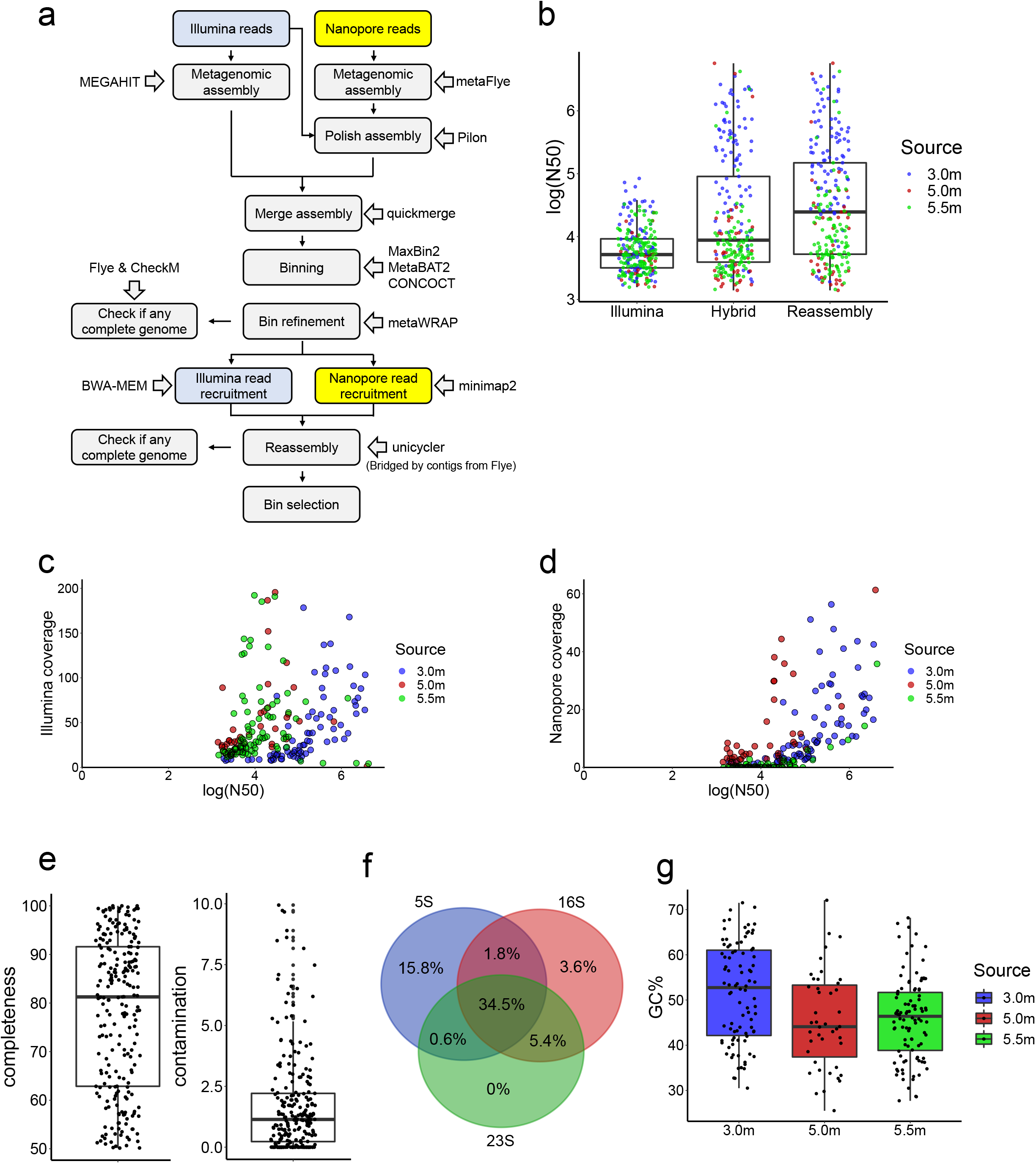
Recovery of genomes from Lake Shunet using long- and short-read sequencing. **a**. The workflow for assembling metagenome-assembled genomes (MAGs). **b**. The value of log (N50) at 3.0, 5.0, and 5.5 m deep using SRs only, combining SRs an LRs (Hybrid), and reassembly of bins using the hybrid method. **c**. The correlation between SR coverage and log (N50) in the final Shunet MAG collection (Reassembly). **d**. The correlation between LR coverage and log (N50) in the final Shunet MAG collection (Reassembly). **e**. The completeness and contamination of recovered MAGs. **f**. Venn diagram from the ratio of MAGs, containing 5S, 16S, and 23S rRNA gene sequences. **g**. The GC ratio of MAGs recovered from the 3.0, 5.0, and 5.5 m deep datasets.

To recover more MAGs and improve contiguity, the assemblies from SRs and LRs were combined before binning. The contiguity of MAGs generated by combining two sequencing reads was dramatically higher than that from the Illumina assembly alone. The average N50 of MAGs from SRs only were 12.4 kb, 6.0 kb, and 7.2 kb in the 3.0, 5.0, and 5.5-m dataset, respectively. Average N50 increased to 476.5 kb, 269.5 kb, and 91.2 kb (Fig. 1b), respectively, when assembling with a combination of the two sequencing methods. A previous study showed that the qualities of MAGs can be improved by reassembly [14], so the step was incorporated into our workflow. When the MAGs were reassembled and selected, the average N50 increased from 476.5 kb to 530.0 kb in the 3.0 m dataset and 91.2 kb to 107.3 kb in the 5.5 m datasets (Fig. 1b).

The correlations between read coverages and contiguity were determined (Fig. 1c, d). The results revealed that the N50 values were more correlated with the Nanopore read coverage (Spearman’s r =0.7) than the Illumina coverage (Spearman’s r= 0.33). This is consistent with the previous observation that contiguity plateaued when the coverage of SRs reached a certain point because the assembly of SRs cannot solve repetitive sequences [9]. Nevertheless, LRs can address the issue by spanning repetitive regions. We also found that using SR assembly only, we cannot obtain MAGs with N50 >100kbp. By comparison, using our workflow, we can obtain 73 MAGs with N50 > 100kbp. The mean SR coverage of these MAGs was 187 times, and mean LR coverage of them was only 67. Additionally, our data size of LRs is about 1/3 that of SRs. Taken together, it represents that the contiguity of MAGs can be greatly improved with one-third LRs. The results highlighted that 1) combining two sequencing methods yield significant improvements in the qualities of MAGs that are recovered from high-complexity metagenomic datasets, and 2) With only extra one-third LRs, we could retrieve genome information, such gene order, from previous SR-derived MAG collections.

Using our workflow, a total of 233 MAGs with completeness > 50% and contamination < 10% were reconstructed. For Genome Taxonomy Database (GTDB) species representatives, the genome quality index, defined as completeness □ -□ 5 □ × □ contamination, should be larger than 50. To meet the GTDB standard, the MAGs were filtered by this criterion, and the MAGs with low SR coverages (<80%) were discarded, resulting in 187 MAGs (Dataset S1). All the MAGs satisfied or surpassed the MIMAG standard for a medium-quality draft [15]. The median completeness of MAGs was 81%, and the median contamination was 1.1% (Fig. 1e). Moreover, 45.3% of the MAGs contained 16S rRNA gene sequences, and 34.5% of MAGs had 23S, 16S, and 5S rRNA gene sequences (Fig. 1f). The median GC ratio of MAGs from 3.0, 5.0, and 5.5 m were 52.75, 44.1, and 46.4%, respectively (Fig. 1g). We also used OPERA-MS to retrieve MAGs from SRs and LRs. However, only 26 medium-quality or high-quality MAGs were recovered, indicating that the method is suboptimal in our case.

### Phylogenetic diversity and novelty of MAGs

To explore the diversity of MAGs, we clustered and de-duplicated the genomes based on a 95% ANI cutoff for bacterial species demarcation [16], since identical microbial species may be detected and assembled from the three different layers. The procedure led to 165 species-level non-redundant MAGs (Dataset S1). The majority (93%) of the species-level MAGs could not be assigned to any known species after taxonomy annotation by the GTDB-Tk, revealing that a great deal of novel MAGs at the species and higher taxonomic ranks were detected (Dataset S2). The novel MAGs comprised six unknown bacterial orders, 20 families, and 66 genera (Fig. 2a).

**Figure 2.**
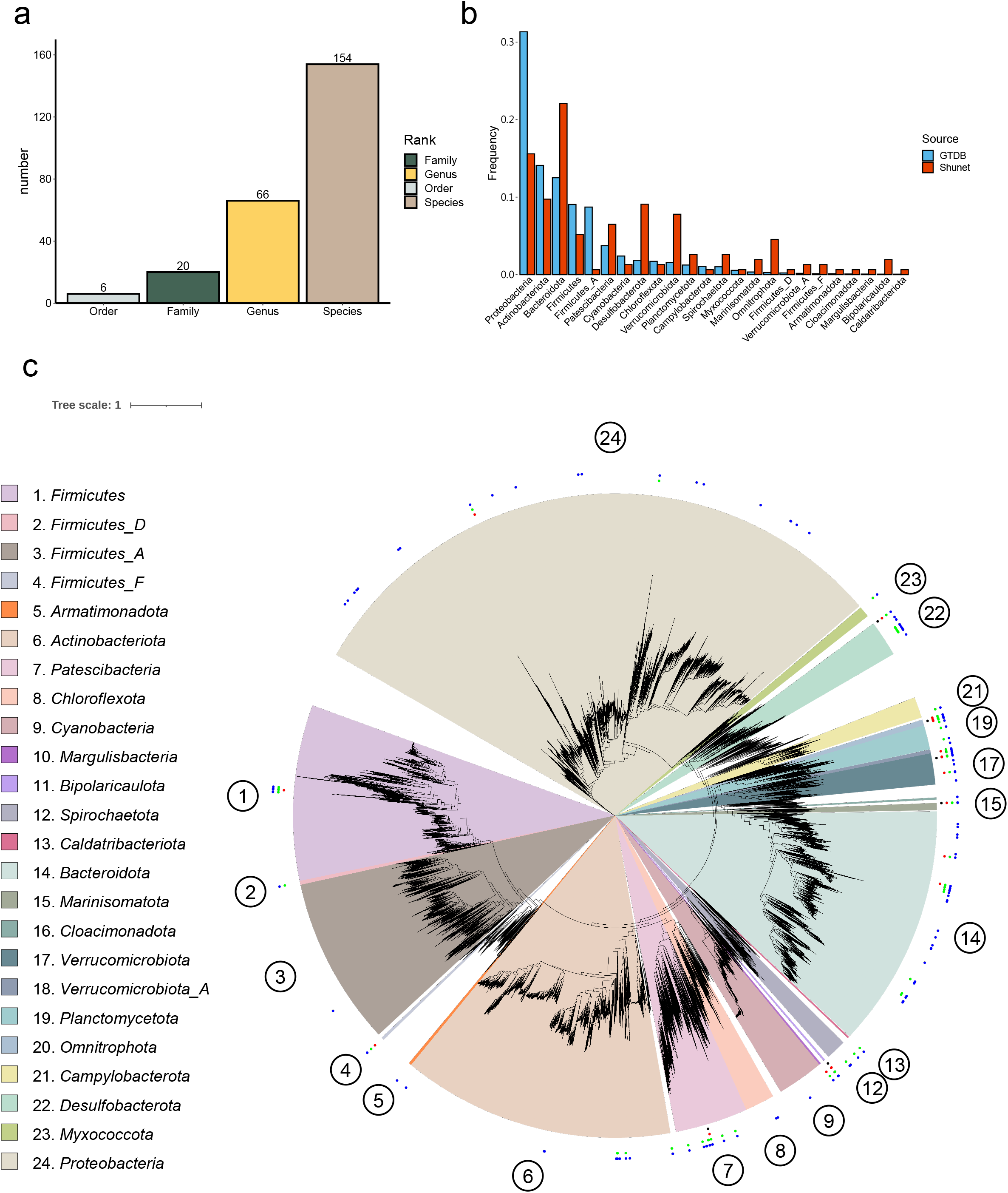
Taxonomical and molecular phylogenetic analyses of recovered bacterial MAGs. **a**. The numbers of novel taxonomic ranks of MAGs assigned by GTDB-Tk. **b**. The phylum frequencies in the MAG collection from the Shunet dataset and GTDB representative genomes. **c**. A phylogenetic tree based on the concatenation of 120 single-copy gene protein sequences. After masking, 5,040 amino acid sites were used in the analysis. The phylogenetic tree includes 188 recovered bacterial MAGs and 30,238 bacterial representative genomes in GTDB-r95. The blue points represent the placement of MAGs that are classified as novel species, the green points represent novel genera, the red points represent novel families, and the black points represent novel orders. Scale bar represents changes per amino acid site.

**Figure 3.**
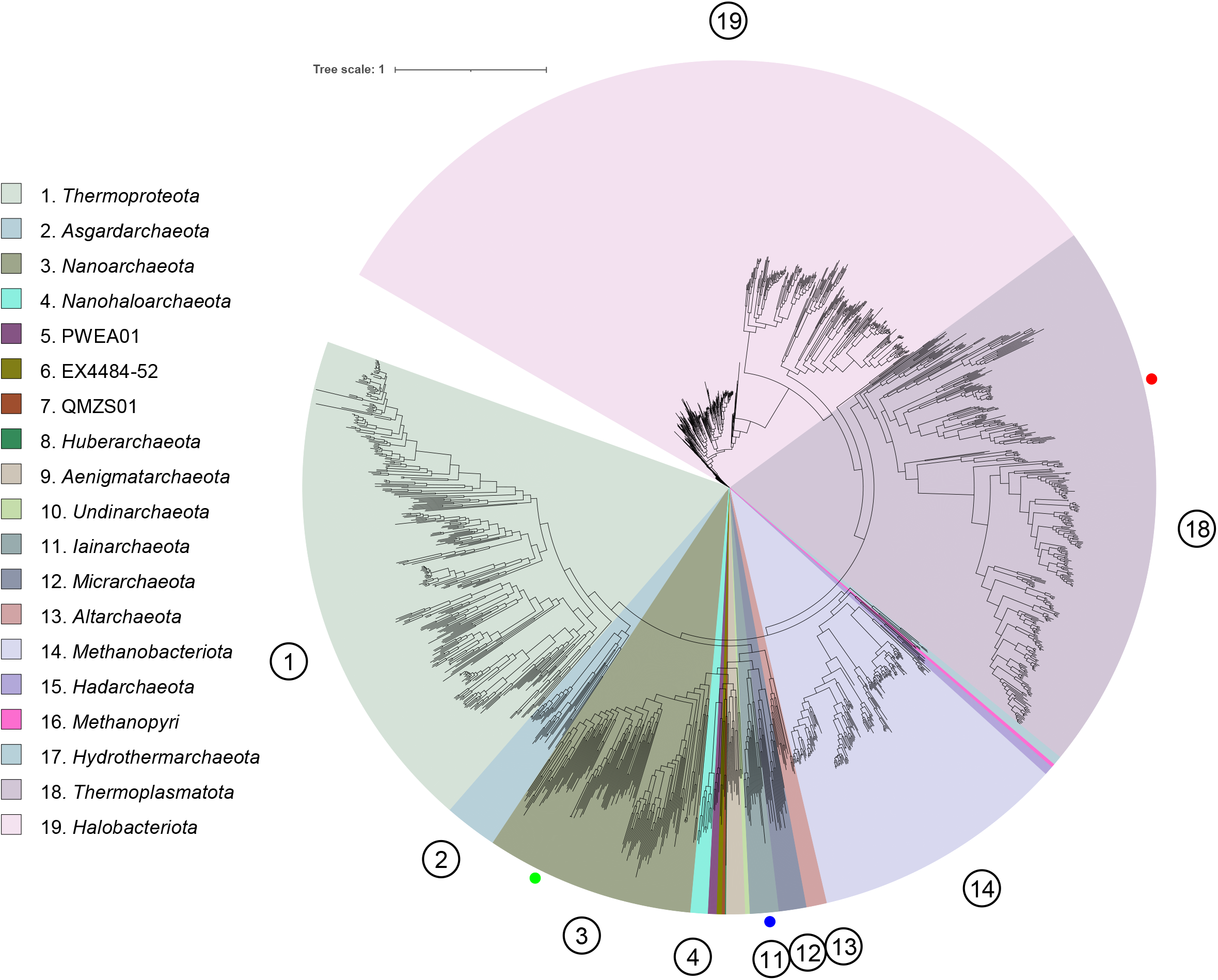
Molecular phylogenetic analysis of recovered archaeal MAGs. The phylogenetic tree was reconstructed based on the concatenation of 122 single-copy gene protein sequences. After masking, 5,124 amino acid sites were used in the analysis. The phylogenetic tree including three MAGs from Lake Shunet and 1,672 archaeal representative genomes in GTDB-r95. The blue dot represents the placement of MAG M55A2, the red dot represents MAG M55A1, and the green dot represents MAG M55A3. Scale bar represents changes per amino acid site.

**Figure 4.**
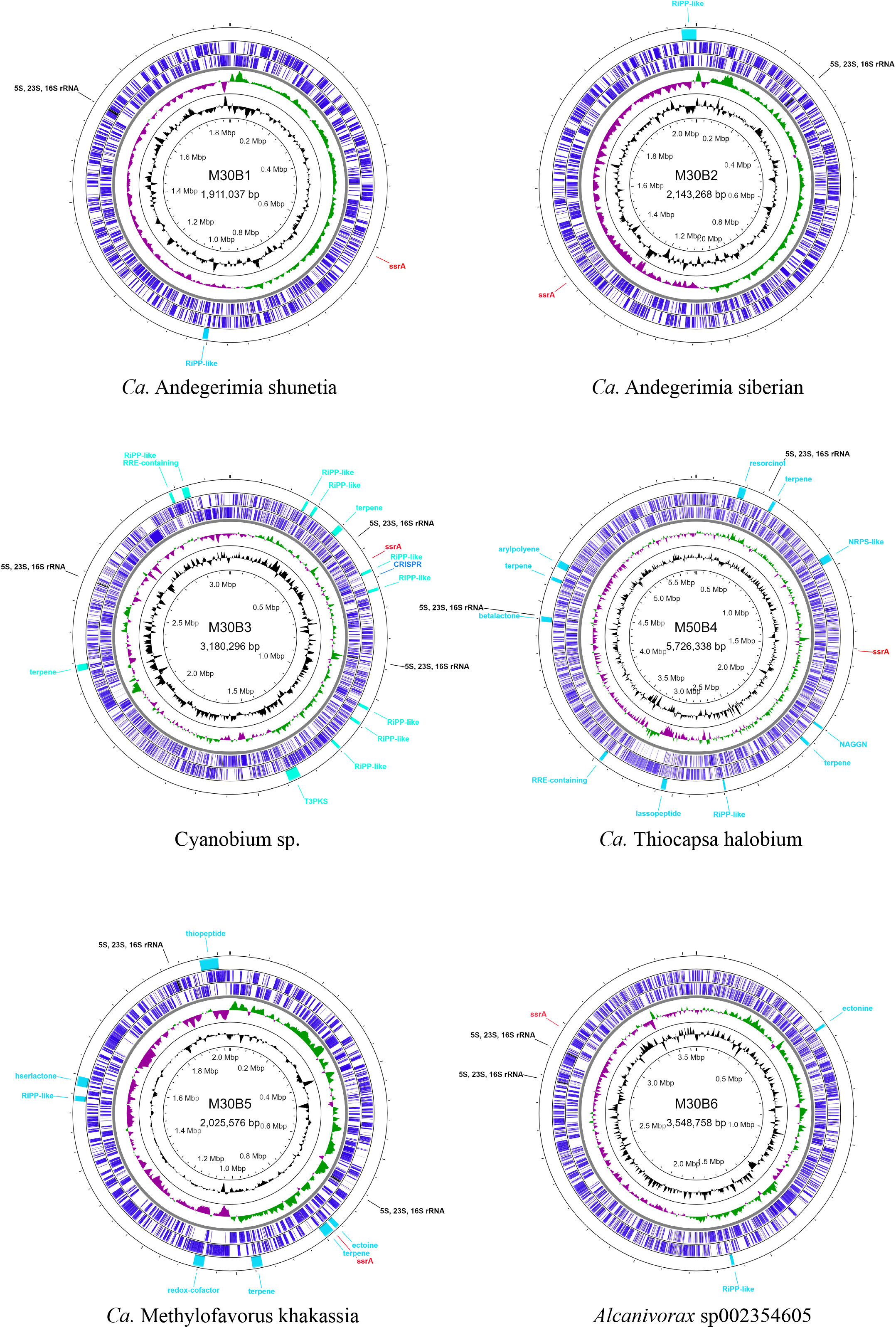
Representation of the six complete MAGs. The rings from the inside to outside represent GC content (black), GC skew-(purple), GC skew + (green), coding sequence regions (blue), rRNA gene sequences (black), transfer-messenger RNA (red), and secondary metabolite gene clusters (light blue). MAG ID M30B6 is classified as *Alcanivorax* sp002354605, M30B1 and M30B2 are *Simkaniaceae* sp., M30B3 is *Cyanobium* sp., M30B5 is “Methylofavorus khakassia”., and M50B4 is “Thiocapsa halobium”.

To examine the phylogenetic diversity in the novel MAGs, a phylogenomic tree was reconstructed using all these bacterial MAGs and representative bacterial genomes in GTDB (Fig. 2b). The result demonstrated that the MAGs widely span the bacterial phylogeny. The MAGs were distributed across 24 phyla, including unusual and poorly-characterized phyla, such as *Armatimonadota, Margulisbacteria, Bipolaricaulota, Cloacimonadota*, and *Caldatribacteriota*. The phylum frequencies differed between the genome collections of the standard database and the Shunet datasets (Fig. 2c). The GTDB mainly comprised *Proteobacteria*. In contrast, in genome collections from the Shunet datasets, the phylum frequency was enriched in the *Desulfobacterota, Verrucomicrobiota, Bacteroidota*, and *Omnitrophota*. The difference can also be seen by comparing Shunet datasets with a genomic catalog of Earth’s microbiomes and 8,000 MAGs recovered by Tyson *et al*. [17, 18], which also had a higher proportion of *Proteobacteria*, but limited in the other four phyla enriched in Lake Shunet. The results suggest that, to gain a comprehensive picture of the microbial genomes on earth, there is a strong need for future studies to explore microbiomes from various habitats, especially overlooked or understudied habitats [17, 19].

### Novel predicted secondary-metabolite biosynthetic clusters and carbohydrate-active enzymes from newly recovered MAGs

Here we demonstrate a) the value of recovering MAGs from rarely investigated habitats to mine novel microbial function potentials and b) the advantage of combining SRs and LRs using two examples: secondary metabolite biosynthetic gene clusters (BGCs) and carbohydrate-active enzymes (CAZymes). Secondary metabolites are usually unique in one or a few species, and not related to the normal growth of the organisms [20]. The secondary metabolites, associated with ecological interactions, can serve as toxins, factors participating in symbiosis with other hosts, defense mechanisms [20, 21]. Identifying novel secondary metabolites enables us to understand the ecological interactions among the microbes. The majority of bacteria remain uncultivated, so mining novel BGCs in metagenomes provides the opportunity to discover new secondary metabolites [22, 23].

In our MAG collection, we identified 414 putative BGCs from 140 MAGs (Fig. S1a). Among them, 134 BGCs were annotated as terpenes and 64 BGCs as bacteriocins. To determine the novelty of these BGCs, the BGCs were searched against the NCBI database using the cutoffs of 75% identity and 80% query coverage based on a previous study [17]. The results demonstrated that 384 BGCs (92%) could not be matched with these thresholds, indicating that most of these could be novel BGCs. Comparably, only 83% of BGCs were predicted to be novel BGCs in the recently-published Genomes from Earth’s Microbiome catalog (GEM) [17].

Complete BGCs are important because they help us identify the metabolites that these BGCs produce using molecular approaches [21]. 72% of BGCs identified from MAGs in the 3.0 m dataset were not on the edge of the contigs, suggesting that the majority of BGCs may be complete. However, only 22% of BGCs in the 5.0 and 5.5 m datasets were not on the edges, which could be because the MAGs from 3.0 m were more contiguous because they had a 10-fold larger median N50 (Fig. 1b). In total, 213 BGCs (51%) we recovered were not on the edges. By comparison, only 34% BGCs predicted in the GEM MAG collection were not on the edge. In the 414 BGCs, 552 core biosynthetic genes, 1,224 additional biosynthetic genes, 205 regulator genes, and 185 transporter genes were identified. This information will enable us to examine the products of the BGC, the function of these genes, and the roles of the products in the individual bacterium. On the other hand, the results also showed that the increased contiguity of MAGs by LRs enables us to obtain more complete BGCs.

Carbohydrate-active enzymes have a range of applications. For instance, CAZymes are used for food processing and food production [24-27]. Exploring novel CAZymes in the metagenome can benefit food industries [24, 25]. On the other hand, identifying novel CAZymes modules enables us to produce novel bioactive oligosaccharides that can be used to develop new drugs and supplements [26, 27]. From the MAGs reconstructed in this study, we identified 8,750 putative CAZymes: 3,918 glycosyltransferases, 3,304 glycoside hydrolases, 738 carbohydrate esterases, and 92 polysaccharide lyases (Fig. S1b). Previous studies indicated that 60∼70% protein identity can be used as a threshold for the conservation of the enzymatic function [28-30]. Among the CAZymes we identified, 1,745 (44%) glycosyltransferases, 1,456 (44%) glycoside hydrolases, 267 (36%) carbohydrate esterases, and 57 (62%) polysaccharide lyases shared less than 60% protein identity with their closest homologs in the NCBI nr database (Fig. S1c). This indicates that these CAZymes could have novel carbohydrate-active functions, which future research efforts can explore further.

### Novel candidate archaeal families identified from Lake Shunet

From the 5.5 m dataset, we identified two MAGs belonging to candidate novel families under *Methanomassiliicoccales* and *Iainarchaeales* (MAG ID: M55A1 and M55A2, respectively) and one MAG belonging to a potential novel species under *Nanoarchaeota*, according to the GTDB taxonomy assignment based on the phylogenomic tree and relative evolutionary divergence (RED) (Dataset S2). In the archaeal phylogenomic tree, M55A1 formed a clade basal to the clade containing species within the *Methanomethylophilaceae* family, a group of host-associated methanogens, and the branch was supported by a 94.7% UFBoot value (probability that a clade is true) [31]. The M55A1 and *Methanomethylophilaceae*-related clade formed a superclade that is adjacent to *Methanomassiliicoccaceae*-related clade, a group of environmental methanogens [32]. These clades formed the order *Methanomassiliicoccales*, the hallmark of which is the ability to produce methane. However, M55A1 did not contain genes encoding for a methane-producing key enzyme complex (Fig. S2). For example, genes encoding methyl-coenzyme M reductase alpha (*mcrA*), beta (*mcrB*), and gamma subunit (*mcrG*), a key enzyme complex involved in methane production, were absent in the M55A1. On the other hand, we did not find *Methanomassiliicoccaceae*-related *mcrA, mcrB*, or *mcrG* genes in the other bins and unbinned sequences in the 5.5 m dataset. Furthermore, M55A1 lacks most of the core methanogenesis marker genes identified in *Methanomassiliicoccales*. The absence of these methanogenesis marker genes implies that the archaea may have lost their methane-producing ability. If this is true, then a phylogenetic group of *Methanomassiliicoccales* may have lost the ability to perform methanogenesis after its ancestor evolved the ability to produce methane. The results not only showed the potential functional diversity in this clade but also highlighted how much such a little-studied environment can reveal about functional diversity in known microbial lineages.

### Five complete MAGs of a candidate novel genus and species from Lake Shunet

The assemblies of Shunet datasets yielded six complete and circulated bacterial genomes. Among these six complete MAG, two belonged to a novel *Simkaniaceae* genus, and there were classified as novel *Cyanobium* species, *Thiocapsa* species, and species under GCA-2401735 (an uncharacterized genus defined previously based on phylogeny), according to the GTDB taxonomy inference based on ANI and phylogenomic analyses (Dataset S1 and S2). The following are individual descriptions of their unique taxonomic and metabolic features. The nitrogen, carbon, sulfur, and energy metabolisms are described in Figure S3.

#### Candidate novel Simkaniaceae genus

According to GTDB-TK, there were two complete MAGs—M30B1 and M30B2—assigned as an unclassified genus under *Simkaniaceae*, a family in the class *Chlamydiia*, based on the topology of the phylogenetic tree. M30B1 and M30B2 formed a monophyletic group and shared 72.48% percentage of conserved protein (PCOP), above the genus boundary of 50% PCOP [33]. The genomes shared 77% ANI, below the 95% cutoff for the same species [16], and the identity of their rRNA gene sequences was 98.45%. Together, the results showed that the two MAGs were two different new species under a novel genus. Therefore, we propose a new genus, *Candidatus* Andegerimia, to include the two MAGs, and renamed the two MAGs as *Candidatus* Andegerimia shunetia M30B1 and *Candidatus* Andegerimia siberian M30B2, abbreviated as M30B1 and M30B2, respectively.

*Simkaniaceae*, like all *Chlamydia*, are obligately intracellular bacteria that live in eukaryotic cells [34]. Validated natural hosts include various multicellular eukaryotic organisms like vertebrates. That some *Simkaniaceae* PCR clones were identified from drinking water implies that *Simkaniaceae* may also live in unicellular eukaryotes [35]. Our samples were collected from saline water, and a membrane was used to filter large organisms. Hence, *Ca*. A. shunetia and *Ca*. A. siberian may be derived from tiny or unicellular eukaryotic organisms.

The reconstruction of complete MAGs enables us to compare genomes in a precise and comprehensive manner by avoiding contamination caused by binning. The two *Simkaniaceae* MAGs we recovered contained five KEGG orthologues that were not present in known *Simkaniaceae* genomes (Table 1). First, the genomes have cold shock protein genes, and the genes were highly conserved (93% amino acid identity) between the two *Simkaniaceae* genomes. Cold shock proteins are used to deal with the sudden drop in temperature [36]. The proteins are thought to be able to bind with nucleic acids to prevent the disruption of mRNA transcription and protein translation caused by the formation of mRNA secondary structures due to low temperature [36]. The existence of the genes in the genomes may confer cold resistance on the *Simkaniaceae* bacteria in Lake Shunet, allowing them to withstand extremely cold environments.

**Table 1.** KEGG orthologues that are present in the novel MAGs but absent in their sister taxa.

|  | **KO number** | **Definition** |
| --- | --- | --- |
| *Simkaniaceae* |  |  |
|  | K00954 | **Pantetheine-phosphate adenylyltransferase** |
|  | K01580 | **glutamate decarboxylase** |
|  | K15736 | **L-2-hydroxyglutarate** |
|  | K01607 | **carboxymuconolactone decarboxylase** |
|  | K03704 | **cold shock protein** |
| *Thiocapsa* |  |  |
|  | K07306 | **anaerobic DMSO reductase subunit A** |
|  | K07307 | **anaerobic DMSO reductase subunit B** |
|  | K01575 | **acetolactate decarboxylase** |
|  | K13730 | **internalin A** |
|  | K05793 | **tellurite resistance protein TerB** |
|  | K05794 | **tellurite resistance protein TerC** |
|  | K05791 | **tellurium resistance protein TerZ** |
| *Cyanobium* |  |  |
|  | K07012 | **CRISPR-associated endonuclease/helicase** |
|  | K07475 | **Cas3** |
|  | K15342 | **CRISP-associated protein Cas1** |
|  | K19046 | **CRISPR system Cascade subunit CasB** |
|  | K19123 | **CRISPR system Cascade subunit CasA** |
|  | K19124 | **CRISPR system Cascade subunit CasC** |
|  | K19125 | **CRISPR system Cascade subunit CasD** |
|  | K19126 | **CRISPR system Cascade subunit CasE** |

Besides the cold shock protein genes, the two *Simkaniaceae* also had glutamate decarboxylase (GAD) genes. GAD is an enzyme that catalyzes the conversion of glutamate into γ*-*aminobutyric acid (GABA) and carbon dioxide. Many bacteria can utilize the GAD system to tolerate acidic stress by consuming protons during a chemical reaction [37]. The system usually accompanies glutamate/GABA antiporters, responsible for coupling the efflux of GABA and influx of glutamate. The antiporter can also be found in the two novel *Simkaniaceae* genomes, indicating that the bacteria can use the system to tolerate acidic environments.

Along with the unique features in the genus, we identified a difference between the two MAGs in terms of metabolism. Taking sulfur metabolism as an example, the M30B2 had all the genes for assimilatory sulfate reduction (ASR), except for *cysH*, and contained the sulfate permease gene (Fig. S3). On the contrary, M30B1 did not contain ASR or the sulfate permease gene. This indicates that M30B2 can take up and use sulfate as a sulfur source, but M30B1 cannot. In summary, the recovery of these genomes broadens our knowledge of the metabolic versatility in *Simkaniaceae*.

#### Candidate novel Cyanobium species

The MAG M30B3 was classified as a novel *Cyanobacteria* species genome under the genus *Cyanobium*, based on phylogenomic tree and 84.28% ANI shared with the Cyanobium_A sp007135755 genome (GCA_007135755.1), its closest phylogenetic neighbor. We named the genome *Candidatus* Cyanobium sp. M30B3, abbreviated as M30B3. The M30B3 is the predominant bacterium in Lake Shunet at 3.0 m and plays a pivotal role in providing organic carbon in the lake ecosystem [12].

Our analysis of the M30B3 genome revealed that the bacterium harbors an anti-phage system that its known relatives lack. In the novel cyanobacterial genome under the *Cyanobium* genus, we found that the genome harbored several CRISPR-associated (Cas) protein genes that were not in other *Cyanobium* genomes (Table 1). The CRISPR-Cas system is a prokaryotic immune system that enables prokaryotic cells to defend against phages [38]. The system can be classified into six types and several subtypes according to protein content. The signature protein of type I is Cas3, which has endonuclease and helicase activities [38]. *cas3* genes can be found in the novel *Cyanobium* genome but not in other known *Cyanobium* genomes. Furthermore, the genome also had *cse1* and *cse2* proteins, signature proteins for the I-E subtype. Our results show that the novel genome harbors a type I-E CRISPR system and that this system is absent in its phylogenetic-close relatives.

#### Candidate novel Thiocapsa species M50B4

Lake Shunet features the extremely dense purple sulfur bacteria (PSB) in its chemocline (5.0 m) layer, and the density of these PSB is comparable to that of Lake Mahoney (Canada), renowned for containing the most purple sulfur bacteria of any lake in the world [39]. A complete MAG of *Thiocapsa* species, the predominant PSB in the 5.0 m layer, was recovered from the 5.0 m dataset. The MAG was classified as a candidate novel species because it shared 90.71% ANI with the genome of *Thiocapsa rosea*. Therefore, we propose the creation of a new species, *Candidatus* Thiocapsa halobium, abbreviated as M50B4.

The complete genome of the predominant PSB M50B4 will help us understand carbon, nitrogen, and sulfur cycling in Lake Shunet. *Thiocapsa* can perform photosynthesis by reducing sulfur as an electron donor, and *Thiocapsa* can fix nitrogen [40, 41]. M50B4 contained genes for bacteriochlorophyll synthesis and the Calvin cycle for carbon fixation. A previous study revealed that *Thioflavicoccus mobilis*, a bacterium close to *Thiocapsa*, can utilize rTCA and the Calvin cycle to fix carbon [42]. In M50B4, all genes for the reverse TCA cycle (rTCA), except for the ATP citrate lyase gene, were identified. Whether the M50B4 can use both rTCA and the Calvin cycle like *T. mobilis* needs to be determined. For sulfur metabolism, the MAG carried intact gene sets involved in SOX system dissimilatory sulfate reduction/oxidation. The sulfate importer gene was also seen in the MAG, which equipped the bacterium with the ability to import extracellular thiosulfate and sulfate and to use them as sulfur sources. In terms of nitrogen metabolism, like other *Thiocapsa*, the bacterium had a gene cluster to conduct nitrogen fixation and a urea transporter and urease gene cluster to utilize urea. Besides nitrogen fixation, currently available *Thiocapsa* have all genes for denitrification, and some *Thiocapsa* have genes to convert nitrite to nitrate. However, the genes were not seen in our MAG.

There are currently five cultured *Thiocapsa* species. Two of these, *T. rosea* and *T. pendens*, contain gas vesicles. Our genomic analysis revealed gas vesicle structure protein genes in M50B4. These genes are also present in *T. rosea* and *T. pendens*, but not in any other *Thiocapsa* genomes, indicating that the genes are critical for vesicles to exist in *Thiocapsa*, and therefore the novel species have gas vesicles. Gas vesicles enable *T*. sp. M50B4 cells to modulate their buoyancy so they can move to the locations with optimal light intensity or oxygen concentration [43]. Environmental conditions of Lake Shunet are known to be dynamic and change with seasons, and this function could be critical for M50B4.

We found that the novel *Thiocapsa* complete MAG have genes that encode dimethyl sulfoxide (DMSO) reductase subunits A and B (Table S2). DMSO reductase is an enzyme that catalyzes the reduction of DMSO into dimethyl sulfide (DMS). The reductase enables bacteria to use DMSO as terminal electron acceptors instead of oxygen during cellular respiration [44]. The DMSO reduction reaction could impact the environment. DMS, the product of the reaction, can be emitted into the atmosphere and be oxidized into sulfuric acid [45]. Sulfuric acid can act as a cloud condensation nucleus and leads to cloud formation, blocking radiation from the sun. The flux of the anti-greenhouse gas DMS is mainly investigated and discussed in oceanic environments [46, 47]. The flux and role of DMS in lake ecosystems are overlooked and rarely documented [48]. Our finding that the extremely dense PSB in Lake Shunet carried DMS metabolism shows the need to investigate the impact and importance of DMS from bacteria in lake ecosystems and sulfur cycling.

#### Candidate novel Methylophilaceae species M30B5

A complete MAG, named M30B5, was classified as a novel *Methylophilaceae* species under a genus-level lineage, called GCA-2401735, which was defined based on phylogenetic placement [49]. The GCA-2401735 lineage currently only comprises two genomes—GCA-2401735 sp006844635 and GCA-2401735 sp002401735—neither of which meet high-quality genome standards due to their low completeness and lack of 16S rRNA gene sequence. The novel complete genome can serve as a representative species of the genus and can be used to infer the capability of the genus (Table S2). Here, we propose the genus *Candidatus* Methylofavorus to include the three GCA-2401735 genomes, and the M30B5 was renamed as *Candidatus* Methylofavorus khakassia.

The isolation locations of the three genomes imply that their habitats were distinct from those of other *Methylophilaceae*. The three “Methylofavorus” genomes were isolated from a cold subseafloor aquifer, shallow marine methane seep, and saline lake, indicating that the bacteria can live in saline environments. By comparison, most other *Methylophilaceae* members live in soil and freshwater or are associated with plants (except for the OM43 lineage) [50]. This indicates that the ancestor of “Methylofavorus” gained the ability to live in saline habitats and diverged from the ancestor of the genus *Methylophilus*, its closest phylogenetic relatives.

The complete genome of M30B5 enables us to comprehensively study metabolic potentials. *Methylophilaceae* is a family of *Proteobacteria* that can use methylamine or methanol as carbon or energy sources [51, 52]. In our analysis, methanol dehydrogenase gene existed in our genome, and methylamine dehydrogenase gene was absent, indicating that the bacteria use methanol as a carbon source instead of methylamine. For motility, flagella are found in some *Methylophilaceae*. Interestingly, flagella- and chemotaxis-related genes were not identified in the MAG but were identified in the other two “Methylofavorus” species, suggesting that M30B5 lacks mobility comparing to the other two “Methylofavorus” species (Fig. S4).

The comparative analysis of M30B5 and other “Methylofavorus” species revealed that the bacteria use different types of machinery to obtain nitrogen (Fig. S4). The formamidase, urease, and urea transporters were present in M30B5 but not the other two “Methylofavorus” species. Instead, the two “Methylofavorus” species had nitrite reductase, which was not in our MAG. The results indicate that M30B5 can convert formamide into ammonia and formate, and take up extracellular urea as a nitrogen source. On the contrary, the other two “Methylofavorus” can use nitrite as nitrogen resources. Our analysis revealed that “Methylofavorus” is metabolically heterogeneous.

## Conclusions

In this study, we successfully developed a workflow to recover MAGs by combining SRs and LRs. This workflow reconstructed hundreds of high-quality and six complete MAGs—including six candidate novel bacterial orders, 20 families, 66 genera, and 154 species—from water samples of Lake Shunet, a meromictic lake with a highly complex microbial community. It demonstrates that with extra less LRs, we can salvage important genome information from previous SR metagenomes. Using comparative genomics, unique and intriguing metabolic features are identified in these complete MAGs, including two predominant novel species: *Thiocapsa* sp, and *Cyanobium* sp. [12]. The findings show that it is advantageous to apply this method in studies of microbial ecology and microbial genomics by revising and improving the shortcomings of SRs-based metagenomes. Additionally, we show that the MAGs contain a high proportion of potential novel BGCs and CAZymes, which can be valuable resources to validate and examine the metabolic flexibility of various microbial lineages through further experimental approaches and comparative genomics. Finally, this study found a high ratio of poorly detectable taxa in the public databases, suggesting that the investigation into rarely explored environments is necessary to populate the genomic encyclopedia of the microbial world, explore microbial metabolic diversity, and fill the missing gaps in microbial evolution.

## Materials and Methods

### Sample collection

Water samples at 3.0, 5.0, and 5.5 m deep were collected from Lake Shunet (54° 25’N, 90° 13’E) on July 21, 2010. The collection procedure was described in our previous research [12]. Briefly, water was pumped from each depth into sterile containers. Part of the water was transferred into sterile 2.0-ml screw tubes (SSIbio^®^) and stored at -80°C until DNA extraction. The rest of the water was filtered through 10-μm plankton and concentrated using tangential flow filtration (TFF) system (Millipore) with 0.22-μm polycarbonate membrane filters. The bacteria in the retentate were then retained on cellulose acetate membranes (0.2 μm pore size; ADVANTEC, Tokyo, Japan) and stored at -80°C until DNA extraction.

### DNA extraction and sequencing

Reads from Illumina and Nanopore sequencing platforms were used in this study. The sequencing reads from Illumina were described in our previous study [12] (Table S1). DNA for Illumina sequencing was extracted from a TFF-concentrated sample using the cetyltrimethylammonium bromide (CTAB) method [53]. In terms of Nanopore sequencing for 3.0-m samples, the same DNA batch used for Illumina sequencing of 3.0-m was sent to Health GeneTech Corp. (Taiwan) for Nanopore sequencing. For 5.0- and 5.5-m samples, there was no DNA remaining after Illumina sequencing, so in 2020 the DNA was extracted again from frozen water samples using the CTAB method by retaining the bacteria on cellulose acetate membranes without TFF concentration. The amounts of DNA were still insufficient for Nanopore sequencing, so the DNA samples were mixed with the DNA of a known bacterium, *Endozoicomonas* isolate, at a 1:2 ratio. No *Endozoicomonas* was detected in the water samples according to our 16S rRNA amplicon survey [12]. The mixed DNA was then sent to the NGS High Throughput Genomics Core at Biodiversity Research Center, Academia Sinica for Nanopore sequencing. To remove reads that had originated from the *Endozoicomonas* isolate, Kaiju web server [54] and Kraken 2 [55] were used to assign the taxonomy for each read; reads that were classified as *Endozoicomonas* by Kaiju or Kraken were removed from our sequencing results. The Nanopore sequencing and processing yielded 13.83, 12.57, and 4.79 Gbp of reads from the 3.0, 5.0, and 5.5 m samples, respectively (Table S1).

### Metagenome assembly

MAG assembly was performed by combining short reads (SRs) from Illumina sequencing and long reads (LRs) from Nanopore sequencing; this workflow is described in Figure 1a. First, the LRs from 3.0, 5.0, and 5.5 m datasets were individually assembled by metaFlye v2.8 [11] with default settings, and the assemblies were polished with corresponding SRs using Pilon v1.23 [56]. On the other hand, SRs were also assembled by MEGAHIT v1.2.9 with k-mer of 21, 31, 41, and 51 [57]. The assemblies from SRs and LRs were then merged by quickmerge v0.3 with parameters -ml 7500 -c 3 -hco 8 [58]. The merge assemblies were then binned using MaxBin2 [59], MetaBAT2 [60], and CONCOCT [61] in metaWRAP v1.3 [14]. The bins from the three bin sets were then refined by the bin refinement module in metaWRAP v1.3. The resulting bins were then polished again by Pilon v1.23 five times. To reassemble the bin, sorted reads that belonged to individual bin were extracted by BWA-MEM v0.7.17 [62] for SRs and by minimap2 for LRs [63]. The extracted long reads were assembled by Flye v2.8, or metaFlye v2.8 if the assembly failed using Flye v2.8 [11, 64]. The bins were then reassembled individually using Unicycler v0.4.8 using the extracted reads and reassembled long-read contigs which were used as bridges [65]. To determine whether the original or reassembled bin was better, the bin with higher value of genome completeness -2.5 × contamination, estimated by checkM v1.1.3 [13], was chosen and retained. Contigs labeled as circular by Flye or metaFlye, >2.0 Mb in size, and completeness >95% were considered “complete” MAGs. The complete MAGs were visualized using CGView Server [66].

While we were preparing this manuscript, Damme *et al*. published a hybrid assembler, called MUFFIN, that also integrates metaFlye and metaWRAP to recover MAGs and Unicycler for reassembly [67]. However, our workflow has a step to merge the assemblies from SRs and LRs to increase the contiguity and assembly size. Moreover, for the reassembly, we use contigs from metaFlye, instead of default setting: miniasm, as the bridge, which we found can produce a better quality reassembly.

### Annotation of metagenome-assembly genomes

The completeness, contamination, and other statistics on metagenome-assembled genomes (MAGs) were evaluated using CheckM v1.1.3 [13]. The genome statistics were processed in R [68] and visualized using the ggplot2 package [69]. The taxonomy of MAGs was inferred by GTDB-Tk v1.3.0 [70]. Average Nucleotide Identities (ANIs) between MAGs were determined by FastANI v1.32 [16]. MAGs were annotated using Prokka v1.14.5 with ‘rfam’ options [71]. To annotate MAGs with KEGG functional orthologs (K numbers), putative protein sequences predicted by Prodigal v2.6.3 [72] were annotated using EnrichM v0.6.0 [73]. The K number annotation results were then used to reconstruct the transporter systems and metabolic pathways using KEGG mapper [74], and the completeness of KEGG modules was evaluated using EnrichM. Secondary metabolite biosynthetic gene clusters in each MAG were identified using antiSMASH v5.0 [75]. Ribosomal RNA sequences were inferred by barrnap v0.9 [76].

### Phylogenetic analysis

Bacterial and archaeal phylogenomic trees were inferred by a *de novo* workflow in GTDB-Tk v1.3.0 [70]. All species-level non-redundant MAGs recovered in this study were analyzed together with the reference genomes in Genome Taxonomy Database (GTDB) [49]. In the *de novo* workflow, marker genes in each genome were identified using HMMER 3.1b2 [77]. Multiple sequence alignments based on the bacterial or archaeal marker sets were then generated and masked with default settings. Trees of bacteria and archaea were then inferred from the masked multiple sequence alignment using FastTree with the WAG+GAMMA models and 1,000 bootstraps [78]. The trees were visualized with the interactive Tree of Life (iTOL) v4 [79].

## Declarations

### Ethics approval and consent to participate

Not applicable

## Consent for publication

Not applicable

## Availability of data and materials

All sequencing data and assembled genomes are available through National Center for Biotechnology Information (NCBI) repositories under BioProject ID: PRJNA721826. Sequence reads of metagenomes from samples at 3.0, 5.0, and 5.5 m deep can be found under SRA accession numbers SRR14300307, SRR14300308, SRR14300309, SRR14307495, SRR14307795, and SRR14307796. The accession numbers of MAGs can be found in dataset S1 and S2.

## Author’s contributions

Y.H.C. and S.L.T. conceived the idea for this study. Y.H.C. assembled the genomes, performed the bioinformatics analysis, and wrote the manuscript. P.W.C. and H.H.C. prepared the DNA samples. D.R. and A.D. collected water samples. S.L.T. supervised the overall study. All authors read and approved the manuscript.

## Acknowledgements

This study was supported by funding from the Ministry of Science and Technology, Taiwan. Y.H.C. would like to acknowledge the Taiwan International Graduate Program (TIGP) for its fellowship towards his graduate studies. We would like to thank Noah Last of Third Draft Editing for his English language editing.

